# Stress-induced collective behavior leads to the formation of multicellular structures and the survival of the unicellular alga Chlamydomonas

**DOI:** 10.1101/2021.08.11.455832

**Authors:** Félix de Carpentier, Alexandre Maes, Christophe H. Marchand, Céline Chung, Cyrielle Durand, Pierre Crozet, Stéphane D. Lemaire, Antoine Danon

## Abstract

Depending on their nature, living organisms use various strategies to adapt to environmental stress conditions. Multicellular organisms implement a set of reactions involving signaling and cooperation between different types of cells. Unicellular organisms on the other hand must activate defense systems, which involve collective behaviors between individual organisms. In the unicellular model alga *Chlamydomonas reinhardtii*, the existence and the function of collective behavior mechanisms in response to stress remain largely unknown. Here we report the discovery of a mechanism of abiotic stress response that Chlamydomonas can trigger to form large multicellular structures that can comprise several thousand cells. We show that these aggregates constitute an effective bulwark within which the cells are efficiently protected from the toxic environment. We have generated the first family of mutants that aggregate spontaneously, the *socializer* mutants (*saz*), of which we describe here in detail *saz1*. We took advantage of the *saz* mutants to implement a large scale multiomics approach that allowed us to show that aggregation is not the result of passive agglutination, but rather genetic reprogramming and substantial modification of the secretome. The reverse genetic analysis we conducted on some of the most promising candidates allowed us to identify the first positive and negative regulators of aggregation and to make hypotheses on how this process is controlled in Chlamydomonas.

## Introduction

When confronted with environmental stress, multicellular organisms implement a set of reactions involving signaling and cooperation between cells from the same tissue. A typical example is the hypersensitive response triggered by plants against pathogens, during which certain cells will be led to self-destruction to enable the entire organism to survive ^1^. In harsh environments, some unicellular organisms such as yeast or bacteria activate defense systems, which also favor the emergence of collective behaviors between cells. These mechanisms involve the formation of multicellular structures such as biofilms ^2,3^. Biofilms, as opposed to aggregates, are surface-anchored structures composed of an extracellular matrix (ECM) made of proteins and sugars, in which cells are embedded to better resist to environmental stresses ^3,4^.

In the unicellular green alga *Chlamydomonas reinhardtii* (hereafter named Chlamydomonas), the formation of multicellular structures has been reported, but the existence and the function of collective behavior mechanisms remain largely unknown. In response to abiotic stress or to predation, Chlamydomonas is able to form clonal assemblies up to 16 cells (palmelloids), or it can passively flocculate by addition of destabilizing agents like multivalent cations or upon pH elevation ^5^. Bigger multicellular structures have been reported in response to predators ^6^, or by selection of sedimenting cells over a long period of time ^7,8^. However, the function of these multicellular structures, their composition, and how their formation is controlled remain to be explored.

Multicellular structures are remarkably interesting from an evolutionary point of view, since they are classically referred to as an intermediate stage in the transition to multicellularity. This is particularly beautifully illustrated in the order Volvocales, which includes unicellular organisms such as Chlamydomonas, life in undifferentiated colonial form, such as *Gonium pectorale*, as well as differentiated multicellular organisms such as *Volvox carteri*^9,10^.

In this study we unravel that the formation of large multicellular structures containing several thousand cells can be induced by a wide range of abiotic stresses, and that this process is a crucial pro-survival mechanism for Chlamydomonas cells. The formation of multicellular structures can be aggregative, when cells assemble together, or clonal when cells do not separate after division ^5^. We demonstrate that in the conditions tested, aggregates are formed by an aggregative process, and that they include a sugar-rich ECM. To understand the genetic basis of this collective behavior, we have generated and screened 13000 insertional mutants, and identified 16 *socializer* mutants (*saz*), which spontaneously form multicellular structures. We characterize here in detail the *saz1* mutant which is defective for the vegetative lytic enzyme (*VLE*) gene ^11^. The aggregates formed by *saz1* result from an aggregative process, and also contain a sugar-rich ECM. Moreover, *saz1* was found to be more resistant to several abiotic stresses. For both *saz1* and stress-induced aggregates, the extracellular compartment had a central role in the formation of multicellular structures. Finally, a large scale multiomics approach taking advantage of the *saz* mutant collection revealed a network of regulators of aggregation in Chlamydomonas.

## Results

### Characterization of aggregation in response to stress

Wild-type cells from the D66 strain (CC4425) ^12^, were subjected to several stresses: S-Nitrosoglutathione (GSNO, nitrosative stress, Morisse et al., 2014), rose bengal (singlet oxygen, Fischer et al., 2005), paraquat (superoxide/H_2_O_2_, Laloi et al., 2007) and heat shock ^16^. For all the stresses tested, at low stress intensity no impact could be visualized in the cultures, while at higher stress intensities cells died. Interestingly, at intermediate stress levels, large aggregates visible to the naked eye were detected and their surface area quantified (Figure 1A). Aggregates could be detected after one day and their size continued to increase steadily with time. To determine whether Chlamydomonas cells form multicellular structures through aggregation or clonal assembly under our conditions, we constructed a strain expressing mVenus, a cytosolic yellow fluorescent protein (YFP). The Venus strain was grown together with the wild type and treated with rose bengal. The multicellular structures obtained were then analyzed and found to be aggregative, since they were composed of mVenus and wild-type cells (Figure 1B). During our observations we found that within the aggregates a translucent structure resembling ECM was very frequently visible (Figure 1C, upper part). Knowing that in bacteria and yeasts the ECM present in biofilms are rich in sugar, we have tried to detect the presence of these compounds in our aggregates. We induced aggregation using rose bengal and stained the cells in the presence of fluorescently-labeled Concanavalin-A (ConA-FITC), that binds glucosyl and mannosyl residues ^17^. In Figure 1C (lower part), it appears clearly that the structures we identified as ECM, exhibit a strong fluorescence showing the presence of sugar. ^2^

**Figure 1.**
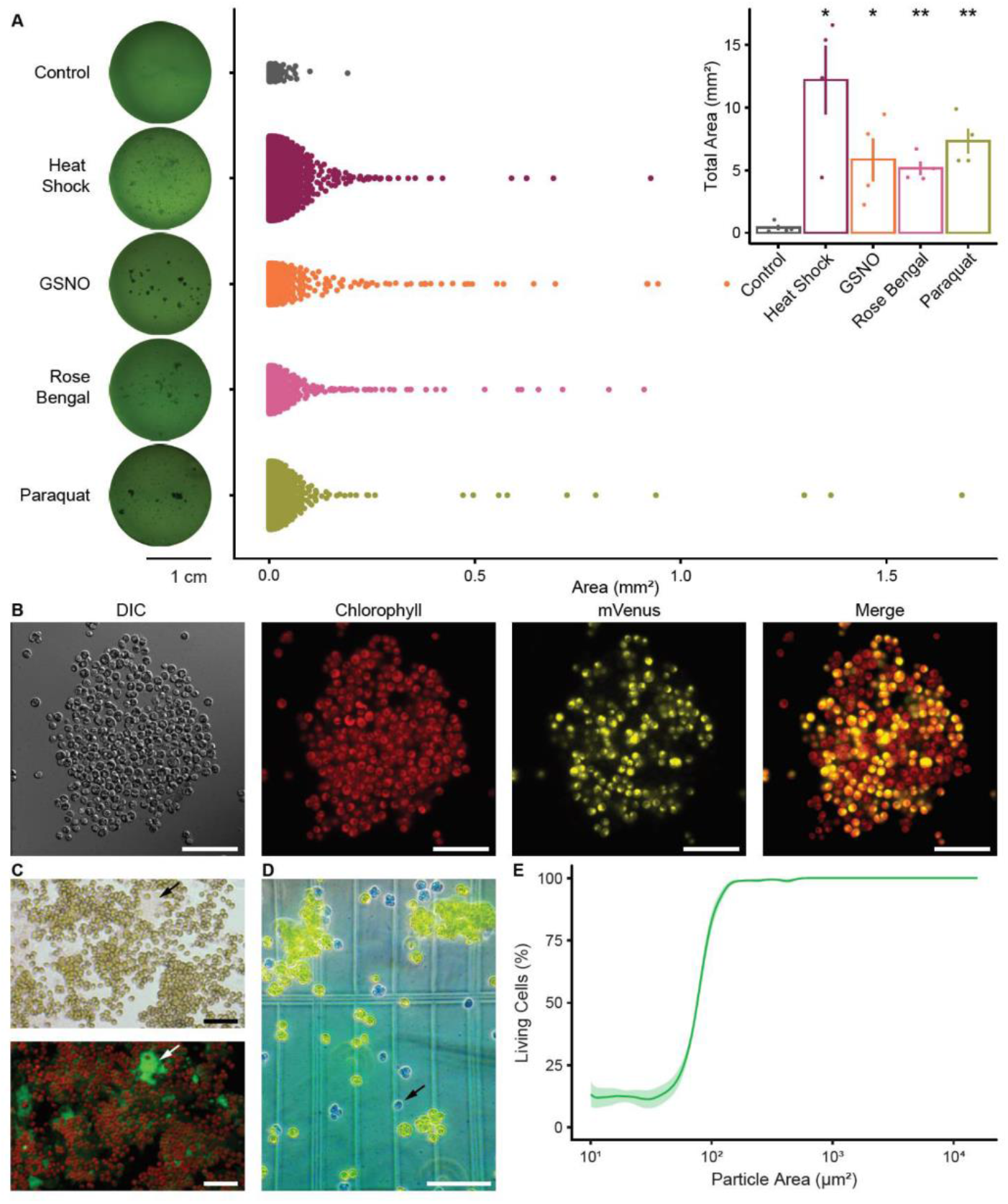
Aggregation in response to stress. (A) Wild-type (CC4425) cultures were exposed to heat shock (50°C, 3 min), GSNO (0.5 mM), rose bengal (2 μM) and paraquat (0.1 μM) and placed in a 24-well plate for 48 h. For each well aggregates were detected, and their surface measured. Each point represents an aggregate and the total of the points represents 4 biological replicates. The histogram represents the average total surface area of the 4 replicates. (B) Microscopic observations of a representative heterogenous aggregate induced by heat shock (50°C, 3 min). Red fluorescence represents chlorophyll and yellow fluorescence represents mVenus. (C) Sugar detection in a representative extracellular matrix (arrow), in an aggregate induced by rose bengal, upper panel imaged in brightfield, lower panel chlorophyll (red) and ConA-FΓΓC fluorescence (green).(D) Labeling of dead cells with Evans blue (arrow) 48 hours after treatment with rose bengal (4 μM).(E) Relationship between the viability of the structures detected and their surface after a 48-hour treatment with rose Bengal (4 μM), “n” represent the number of objects analyzed from four biological replicates. Error bars (A) and shaded area (D) indicates ±SEM and for *t*-test: * p <0.05 or **p ≤ 0.01. Scale bars represents 50 μm (B and E) and 100 μm (C).

To test whether the formation of aggregates could enable increased stress resistance, we treated wild-type cultures with rose bengal, and stained dead cells using Evans blue ^18^. We observed that most of the time, within multicellular structures, cells were protected from the toxic environment, while the others were dead (Figure 1D). To quantify this phenomenon, we developed an ImageJ ^19^ macro allowing detection of all the structures, their size and whether the cells are alive (green) or dead (blue). These analyses clearly showed that the larger the structures to which the cells belong, the more the phenomenon of protection is exacerbated (Figure 1E).

### *Socializer* 1 mutant

To understand the genetic mechanisms that control the process of aggregation, we created and screened a libr ary of 13,000 insertional transformants. This allowed identification of 16 mutants dubbed *socializer* (*saz*) for their ability to aggregate spontaneously. In the present study we have extensively characterized the mutant *saz1* which forms large multicellular structures visible to the naked eye and containing several thousand cells (Figure 2A). The size of *saz1* aggregates varied between 100 μm^2^ and several mm^2^, in the same range as stress-induced aggregates of wild type cells (Figure 2C). When observed under the microscope, we saw that the cells form a homogeneous structure in which they appear to be attached to each other (Figure 2D). The *SAZ1* locus was characterized using the protocol described previously ^20^. The *saz1* mutant harbors an insertion in the Cre01.g049950 gene (Figure 2B), previously named vegetative lytic enzyme (*VLE*)^21^ or sporangin (*SPO*) ^11^. We precisely localized the insertion in the exon 21 of Cre01.g049950 and verified that a wild-type copy of the corresponding mRNA was no longer expressed in *saz1* (Figure S1). To confirm that a mutation in the *VLE* gene leads to a spontaneous aggregation phenotype, we analyzed three additional insertional *vle* mutants obtained from the CLiP library ^22^. In these mutants with an insertion respectively in the 1^st^ exon, the 6^th^ exon and the 17^th^ intron, the *VLE* mRNA could not be detected (Figure S1). Similarly to *saz1*, these three mutants aggregated spontaneously as opposed to the corresponding wild-type strain (CC5325) (Figure 2C).

**Figure 2.**
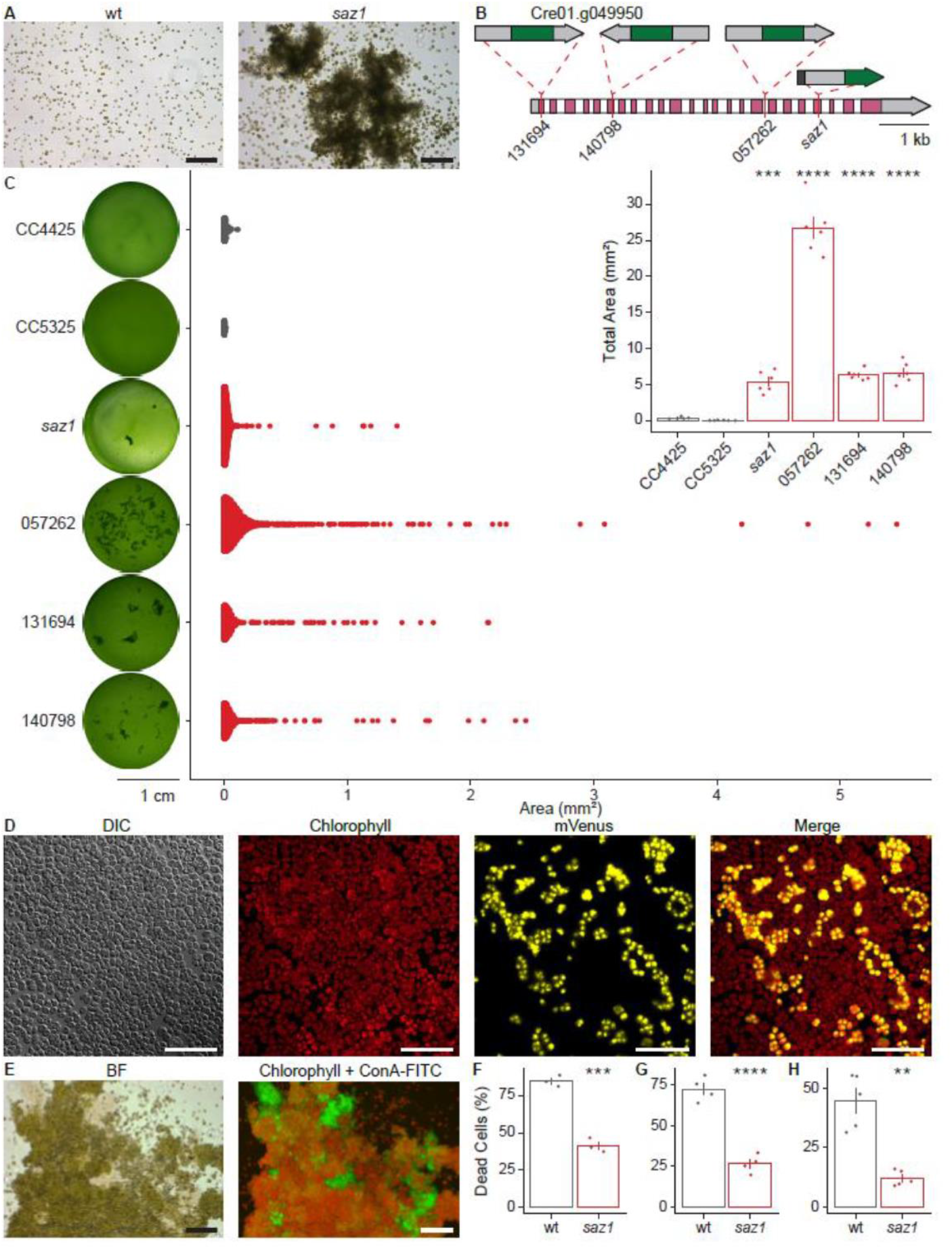
*Socializer1* characterization. (A) Representative microscopic observation of cultures of the wild type (CC4425) and *saz1* mutant under standard conditions. (B) Insertion sites of *saz1* and three CLiP mutants (LMJ.RY0402.<u>131694</u>. LMJ.RY0402.<u>140798</u> and LMJ.RY0402.<u>057262</u>) in the *VLE* gene (Cre01.g049950). (C) Analysis of *saz1*, the CLiP mutants and the corresponding wild-types (CC4425 and CC5325), in 24 well plate. For each well aggregates were detected, and their surface measured. Each point represents an aggregate and the total of the points represents 6 biological replicates. The histogram represents the average total surface area of the 6 replicates. (D) Microscopic observations of a representative aggregate formed in a co-culture of wild-type and Venus (YFP) strains. Red fluorescence represents chlorophyll and yellow fluorescence represents mVenus. (E) Sugar detection in a representative extracellular matrix (arrow), within a *saz1* aggregate, left panel brightfield, right panel chlorophyll (red) and ConA-FITC fluorescence (green). (F) Death quantification in wild-type and *saz1*, after (F) heat shock (50°C - 4 min. 24 h), (G) GSNO (1 mM, 24 h) and (H) rose bengal (4 μM, 8 h) treatments.Error bars indicates ± SEM and for *t*-test: * p ≤ 0.05 or ** p ≤ 0.01. Scale bars represents 50 μm (D) and 100 μm (A, H).

To determine the way aggregates are formed in *saz1*, we grew cultures composed of a combination of *saz1* and the mVenus strain described in Figure 1B. As in the case of the aggregates formed in response to stress by wild-type cells, the multicellular structures spontaneously formed by *saz1* were heterogenous and therefore the result of an aggregative process (Figure 2D). A sugar-rich ECM could be detected in *saz1* aggregates using ConA-FITC (Figure 2E). The formation of wild-type aggregates allowed increased resistance of the cells to stress conditions (Figure 1C-D). Consistently, *saz1*, which forms aggregates spontaneously, was also more resistant to stress. Indeed, *saz1* showed a higher resistance to all the stresses tested (GSNO, rose Bengal and heat shock), (Figure 2F-H), confirming the protective role of aggregates.

### The culture medium is sufficient to induce aggregation

Growing *saz1* with the mVenus strain that does not spontaneously aggregate, induces the formation of heterogenous multicellular structures (Figure 2D), suggesting a potential role of the culture medium in transmitting an aggregation signal. To test this hypothesis, we removed the *saz1* cells from the culture medium by centrifugation and filtration. Wild-type cells were then grown in this culture medium, and the formation of multicellular structures was quantified. Remarkably, the culture medium of the mutant was able to induce a strong aggregation of wild-type cells (Figure 3A). There is therefore probably one or several molecules in the culture medium capable of inducing aggregation. To investigate the nature of this molecule we subjected the culture medium to different treatments. After boiling for two minutes, a treatment that denatures proteins, the culture medium was no longer able to induce aggregation (Figure 3A). Filtration of the culture medium revealed that only the fraction containing molecules above 30 kDa was capable of inducing aggregation (Figure 3A). These results are consistent with the hypothesis that one or several proteins in the culture medium might induce aggregation. To test whether the culture medium resulting from stress-induced aggregation could also induce the formation of aggregates, we induced aggregation using heat shock, as this treatment does not require incorporation of foreign molecules into the culture medium. This culture medium was just as capable as the medium of the *saz1* mutant to trigger aggregation of non-stressed cells (Figure 3B).

**Figure 3.**
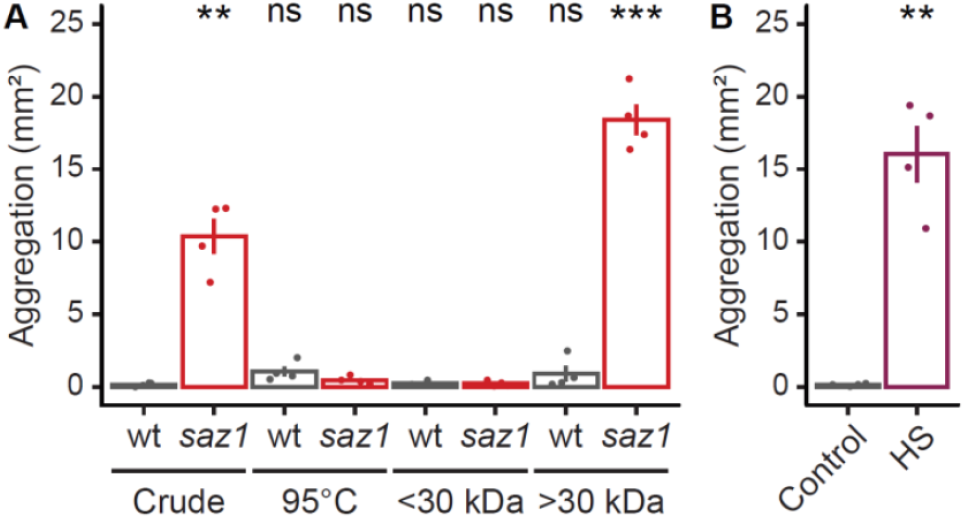
Induction of aggregation by the culture medium. (A) Medium from wild-type (CC4425) and *saz1* cultures were harvested and used to begin new wild-type cultures in 24-well plates. Other cultures were initiated in wild-type (grey) or *saz1* (red) medium that were boiled for 2 minutes or fractionated using a 30 kDa filter. For each culture, aggregates were detected, and their surface measured. Values represent the average of total surface area of 4 independent wells. (B) Wild-type cells were grown in 24-well plates in control (grey), or heat shock (purple) medium. Values represent the average of total surface area of the aggregates detected in 4 independent wells.Error bars indicates ± SEM and for the *t*-test * p ≤ 0.05; ** p ≤ 0.01: *** p ≤ 0.001 or **** p < 0.0001.

### The aggregation secretome

As the culture medium is sufficient to induce aggregation in Chlamydomonas, possibly by way of proteins, we performed a quantitative proteomic analysis by comparing the culture media of 6 of the *saz* mutants having the most pronounced phenotypes, with that of the wild type. For each strain, five biological replicates were prepared and analyzed by tandem mass spectrometry. In our seven samples we identified a total of 763 proteins in the culture medium (Table S1). The PredAlgo algorithm ^23^, predicts that 44.7% of these proteins are secreted, 14.2% are chloroplastic, 5.8% are mitochondrial while 36.3% of them have no prediction. Consistent with the fact that the culture medium was analyzed, a majority of the identified proteins are predicted to be secreted. The proteins whose content is altered in several mutants in comparison to the wild type, were classified into different categories, depending on whether their expression is altered in 6, 5, 4, 3 or 2 mutants (Figure 4A and B). To identify common regulators of aggregation we focused on the proteins whose content is significantly deregulated in 6 but also in 5 mutants (131 proteins), to avoid missing interesting candidates. We classified the proteins into different categories according to their annotation on the JGI Phytozome v13.0 website (https://phytozome-next.jgi.doe.gov), and their descriptions in the literature (Figure 4C).

**Figure 4.**
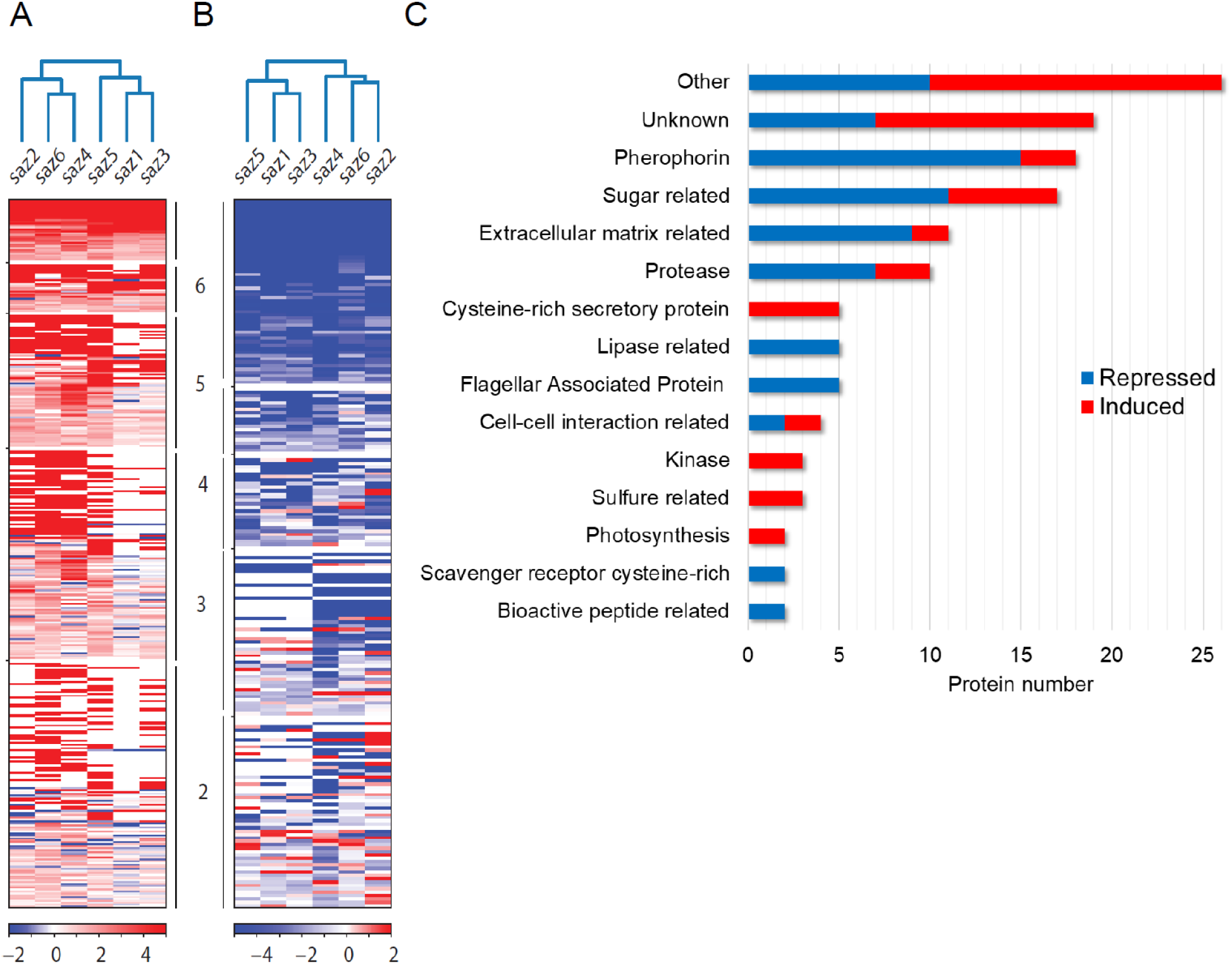
Quantitative proteomic analysis of the extracellular medium. We analyzed the protein content of the culture medium in the wild type in comparison to 6 different *saz* mutants. The results are expressed as a ratio of the different mutants compared to the wild type, we classified them according to whether the proteins are under-represented (A) or over-represented (B) in 6, 5, 4, 3 or 2 mutants. (C) Function of the proteins deregulated in 5 or 6 mutants according to the number of proteins represented.

In this list pherophorins hold an especially important place (16/131). These proteins are essential constituents of the ECM in *Volvox carteri*, a multicellular organism that belongs, like Chlamydomonas species, to the order of Volvocales ^24^. Among the most represented protein classes, we also found proteases, including the subtilisin VLE which is absent in *saz1* as expected, and interestingly downregulated in the medium of four other *saz* mutants. We also detected several matrix metalloproteinases (MMPs), such as gametolysins which are known to degrade the cell wall during mating in Chlamydomonas ^25,26^, but also to contribute to the destruction of the ECM in *Volvox carteri* ^27^ and animals ^28^. Other proteins related to the ECM were also detected, such as VSPs (vegetative, SP rich) ^29^, lysyl oxidases (LOX) containing scavenger receptor cysteine-rich (SRCR) domains, which are necessary for ECM formation in animals ^30^, or formin which influences ECM stability through the control of actin polymerization ^31,32^. We also found many sugar-related proteins, mainly involved in catabolism (e.g., Glucan 1,3-beta-glucosidase or Beta-galactosidase), glycosylation or sugar binding (lectins). Interestingly, we found four ephrin A/B receptor-like proteins that are known in animals to mediate cell-cell adhesion ^33^. Two of them share similarities with mastigonemes which have been proposed to play a role in adhesion through an interaction with the transient receptor potential cation channel PKD2 (polycystic kidney disease 2) ^34^. The latter mediates flagellar agglutination during mating ^35^. Ephrin-related genes have also recently been identified during the formation of multicellular structures in response to exposure to a predator ^36^. Altogether this suggests that adhesion mechanisms could also have a role in the assembly of multicellular structures. Finally, we found several lipases that are downregulated in *saz* mutants, suggesting that lipids may be important for the aggregation process (Figure 4C). The entire list of the 131 proteins is shown in Table S1.

### Transcriptomic analysis of the *saz* mutants

We completed our systemic approach by analyzing the transcriptome of the same six *saz* mutants in comparison with the wild type, using mRNA Illumina High-Sequencing technology. During this analysis, a total of 16,829 transcripts were detected and quantified, (Table S2). As we did for our proteomics analysis, we focused on genes deregulated in 6 but also in 5 mutants, to avoid missing important genes that would not be affected in one mutant (Figure 5 A and B). This subset corresponds to the 249 genes significantly up- or down-regulated at least 4-fold in at least 5 *saz* mutants (Table S2).

**Figure 5.**
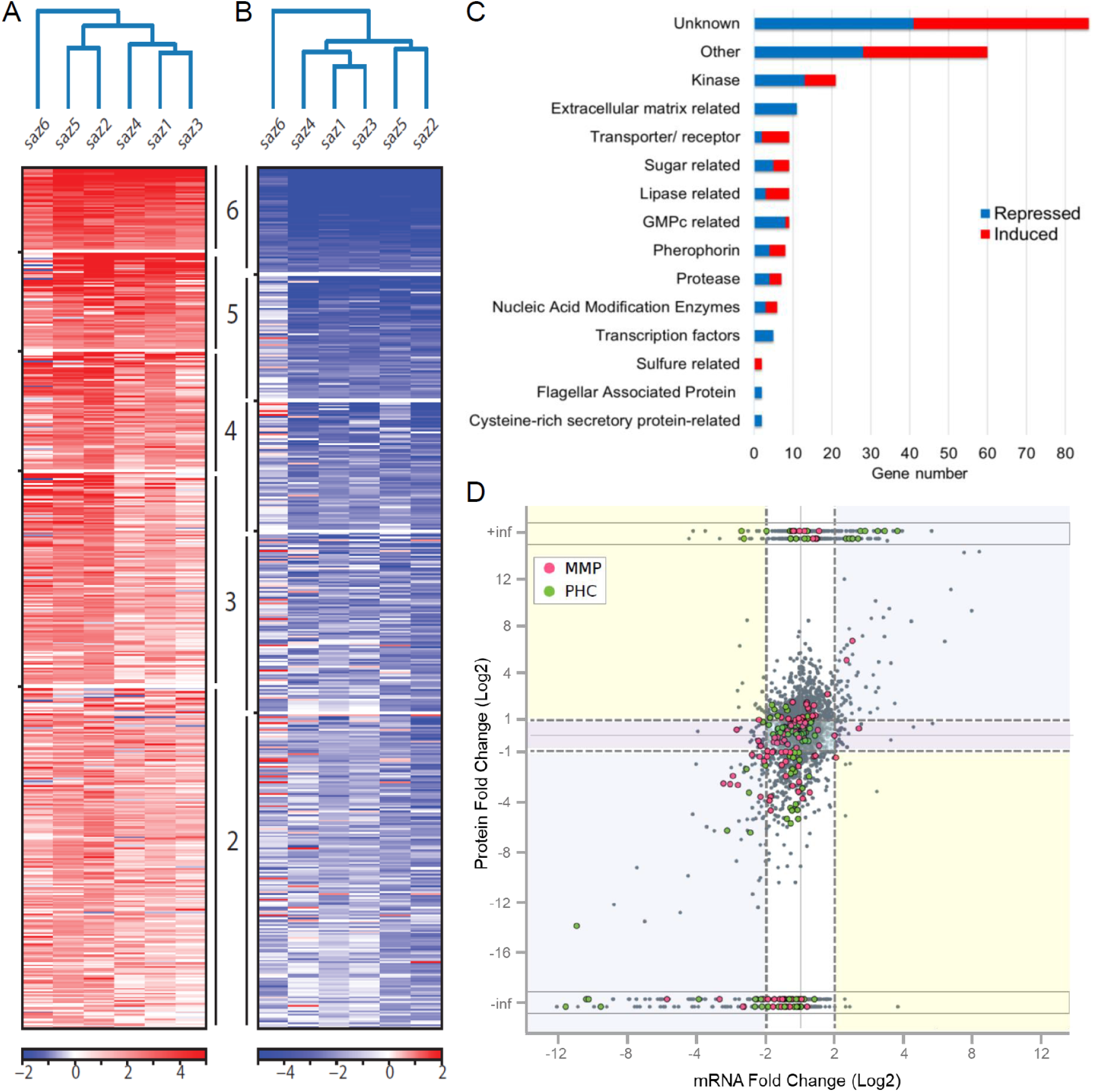
Transcriptomic analysis of the *saz* mutants. We analyzed the transcriptome of the wild type in comparison to 6 different *saz* mutants. The results are expressed as a ratio of the different mutants compared to the wild type, we classified them according to whether the genes are repressed (A) or induced (B) in 6, 5, 4, 3 or 2 mutants. (C) We analyzed the function of the genes that are deregulated in 5 or 6 mutants and we divided them into different categories according to the number of gene represented. (D) Cross-analysis of the proteomic and the transcriptomic data. We compiled the results obtained during our proteomics and transcriptomics studies for the 747 proteins whose gene expression could also be quantified. Each of the points reflects the ratio between a mutant and the wild-type strain. When a protein has been found in a mutant and not in the wild type or vice versa, an infinite value is assigned to it (-inf or +inf, in grey rectangles). When the regulation is done in the same way at the protein and gene expression level the dots are in a blue rectangle, if the regulation is done only at the protein level the dots are in a white rectangle, if the regulation is done only at the gene expression level the dots are in a purple rectangle and if the gene expression and proteomic regulations go in opposite directions the dots are in a yellow rectangle. We have highlighted the example of pherophorins (PHC). which appear as green dots, and matrix metalloproteinases (MMP). which appear as pink dots.

Strikingly, many genes belong to the main families identified during our proteomic analysis including several genes encoding pherophorins, proteases, lipases, as well as proteins linked to the ECM (Figure 5C). We also identified several genes encoding lectins, that bind specifically to sugars. Beyond these families also found in our secretome, this transcriptomic analysis uncovered additional gene families that may be involved in the intracellular signaling controlling aggregation. Sixteen genes encode kinases, which are key players in cellular signaling. Interestingly, six genes related to cyclic guanosine monophosphate (cGMP), which can act as a second messenger controlling kinase activities ^37^, are downregulated in *saz* mutants, including four guanylate cyclases. Other interesting genes with a potential role in sensing or signaling include several ion channels, receptors, or transporters (Figure 5C).

### Cross-analysis of the transcriptomic and proteomic analyses

Of the 763 proteins identified in the aggregation secretome, 747 of the corresponding mRNA were quantified in the transcriptomics study. We studied the regulation of these 747 proteins in comparison to the expression of the corresponding genes, in the six *saz* mutants compared to the wild type (Figure 5D). Strikingly, we observed that the variation in the amount of protein in the extracellular compartment is most often not accompanied by a variation in the expression of the corresponding gene. Indeed 56.9% of proteins are present in different quantities in the extracellular compartment without being affected in the expression of the corresponding gene (Figure 5D, white rectangles), whereas only in 36.9% of cases the same behavior is observed at the gene and protein levels (Figure 5D, blue rectangles). In very few cases (3.7%) we observe a deregulation of gene expression without the corresponding protein being affected (Figure 5D, purple rectangles) and in even rarer cases (2.5%), the regulations at the level of gene expression and the corresponding protein go in opposite directions (Figure 5D, yellow rectangles). These results indicate that other mechanisms than transcription are most often responsible for the changes in protein abundance in the culture medium such as changes in mRNA or protein stability, in translation or subcellular localization. To illustrate this phenomenon in further details, we focused on the pherophorins and the MMPs that were detected in the extracellular compartment (Figure 5D, green and pink dots respectively). For many members of these two families, we find both under-expressed and over-expressed representatives, illustrating the complexity of the signaling pathways induced during aggregation, and that these protein families could contain members that are pro-aggregative as well as anti-aggregative.

### Identification of new regulators of aggregation

Our multiomic analysis allowed identification of candidates potentially involved in the control of aggregation. To assess this possible function, we analyzed insertion mutants of the CLiP library ^22^, for some of the genes of the main categories identified. We selected several pherophorins, MMPs and VSPs from our proteomic or transcriptomic analyses. Among the selected mutants, several exhibit a phenotype related to aggregation, which we have characterized in more detail. For each mutant strain, we confirmed the presence of the insertion sequence at the expected locus identified by the CLiP library (Figure S2). The ability of each mutant to aggregate in response to stress was investigated. Three mutants affected in pherophorin genes (*PHC30, PHC41* and *PHC50*), which are all up regulated in *saz* mutants, are no longer able to aggregate in response to rose bengal (Figure 6A). This suggests that the corresponding proteins have a pro-aggregation role. Conversely, several mutants spontaneously form aggregates, indicating that the corresponding proteins probably have an anti-aggregation role (Figure 6B). For all these mutants, the anti-aggregation role is consistent with the downregulation of the corresponding protein (PHC28, PHC35, MMP32, VSP4) or mRNA (PHC28, PHC35) revealed by our multiomic analysis (Figure 6B).

**Figure 6.**
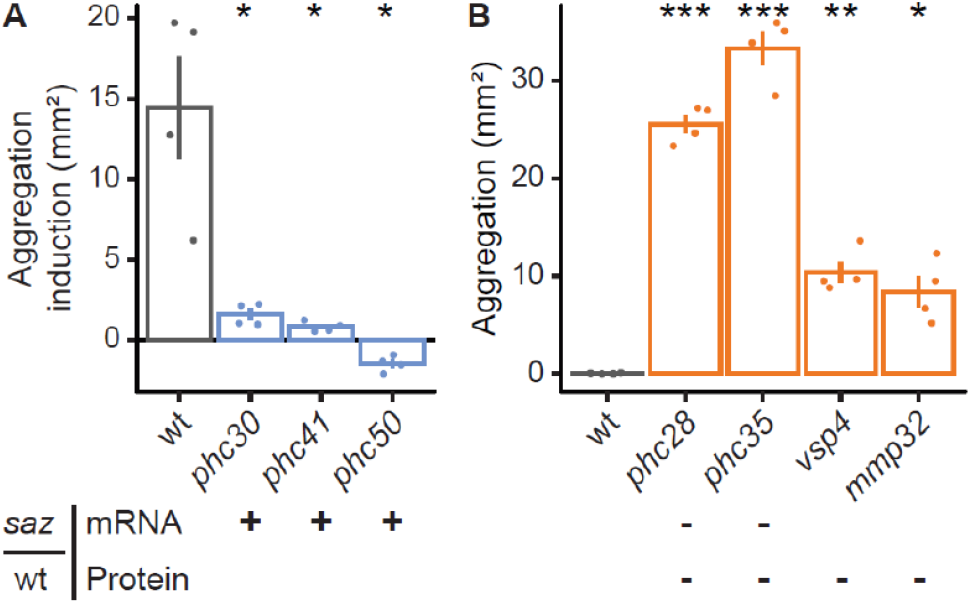
Reverse genetic analysis. From our multiomics analysis, we have selected and characterized CLiP mutants of genes of potential interest, strains exhibiting an aggregation-related phenotype are presented here. These strains were grown in 24-well plates and aggregates were detected and their surface measured. For each condition, values represent the average of total surface area of 4 biological replicates. Some of the mutants we analyzed are no longer able to form aggregates in response to rose bengal (A), other mutants spontaneously form aggregates under standard culture conditions (B). When a gene or protein is down- or up-regulated in *saz* mutants, the symbols (-) or (+) is notified under its name.

## Discussion

Several responses to abiotic stress have been reported in Chlamydomonas including acclimation, palmelloid formation and programmed cell death ^5^. Here we report the discovery of a mechanism of abiotic stress response involving collective behavior between Chlamydomonas cells and involving the formation of large multicellular structures visible to the naked eye, that can comprise several thousand cells (Figure 1A). This phenomenon appears as a general stress response since aggregation is induced and can confer resistance to stresses of different nature. The mode of aggregate formation is not simply the result of the non-separation of cells after division (clonal mode), but rather a complex process that may involve communication between cells, a change in their ability to interact and a sugar-rich ECM (Figure 1B-C). We show here that the formation of aggregates is an efficient way for Chlamydomonas to survive abiotic stress, and that the size of aggregates correlates with the capacity to survive under stress conditions (Figure 1D-E).

To understand how this complex process is genetically controlled, we have identified the first family of mutants exhibiting a spontaneous aggregation phenotype, the socializer (*saz*) mutants. A large scale multiomic analysis of six of these mutants revealed the existence of a common genetic program and the importance of the protein composition of the extracellular compartment to implement a collective behavior leading to the formation of multicellular structures in response to stress. Spontaneous aggregation of *saz* mutants appears comparable to stress-induced aggregation including the same mode of aggregate formation, a sugar-rich ECM within multicellular structures (Figure 2D-E), and increased resistance to stress (Figure 2D-E). The *saz* mutant collection therefore constitutes an invaluable tool for the understanding of this phenomenon.

In the present study we further characterized the *saz1* mutant and showed that inactivation of the Cre01.g049950 gene induces a spontaneous aggregation phenotype (Figure 2A-C). This gene has already been described in the literature as vegetative lytic enzyme (*VLE*)^21^ or sporangin (*SPO*) ^11^. Because of its expression profile, its spatio-temporal localization and its biochemical function, it was suggested that VLE was responsible for sporangial cell wall degradation during cell division ^11,21^. However, this hypothesis has never been validated by the characterization of a VLE mutant. If the sole function of VLE was to release cells during division, a culture of the *vle* mutant would likely consist only of cells in sporangia or possibly in larger multicellular structures resulting from a clonal formation process. By contrast, for the four *vle* mutants analyzed, we observed both free cells, as well as large aggregates (Figure 2A), which we showed to be heterogeneous and therefore not the result of a clonal formation process (Figure 2D). This would suggest that not only VLE, but several enzymes are capable of releasing cells during division, and that VLE has at least one other function in controlling aggregation in Chlamydomonas. Other authors have suggested that VLE could be a gametolysin activator ^38^. Gametolysins are MMPs that are responsible for cell wall removal during mating and need to be activated by a still unknown serine protease ^39^. Therefore VLE, which was the only serine endoprotease identified in the mating secretome ^38^, was proposed to be responsible for the cleavage activation of gametolysins during this process. MMPs are also known to control the degradation of the ECM in plants ^40^ and animals ^28,41^. The possible MMP activation function of VLE could explain how it contributes to the control of aggregation in Chlamydomonas, as ECMs are present in all the aggregates we analyzed (Figure 2E). Another argument in favor of the importance of MMPs in the control of the aggregation process is that several of them are deregulated in *saz* mutants, at the gene and/or the protein level (Figure 5D). To investigate the potential involvement of the selected MMPs, the corresponding mutants were characterized. A mutant of MMP32 exhibits a spontaneous aggregation phenotype, giving it an anti-aggregation role in this process (Figure 6B). Beyond MMPs which are known to target ECM proteins ^40,41^, we identified other protein controlling ECM formation in other organisms. Pherophorins are a family of extracellular hydroxyproline rich proteins, whose function remains poorly known in Chlamydomonas. In *Volvox carteri*, a multicellular green alga belonging like Chlamydomonas to the order of Volvocales, pherophorins are known to be necessary for the formation of the ECM ^24^. A pherophorin was also proposed to participate in the signaling process leading to sexual reproduction in *Volvox carteri* ^42^. In both our transcriptomic and proteomic analyses, we found many pherophorins which are over- or underexpresed in *saz* mutants. The role of phrerophorins in relation to aggregation in Chlamydomonas appears complex, since while some pherophorins are present in all *saz* mutants and absent in the wild type, which one would expect from a protein necessary for the formation of the ECM, the opposite situation is also true and even more frequent (Figure 5D). There would thus exist pherophorins having a pro-aggregative or anti-aggregative role. This hypothesis has been confirmed by our reverse genetic analysis, which shows that when pherophorins are absent in *saz* mutants and present in the wild type (PHC28, PHC35), the corresponding mutants spontaneously aggregate, whereas when they are absent in the mutants and present in the wild type (PHC30, PHC41, PHC50), their mutants are no longer able to aggregate in response to stress (Figure 6A). Pherophorins have A and B globular domains similar to plant lectins, suggesting that beside being glycosylated, they may also have sugar-binding functions ^24^, such as an interaction with the sugar-rich ECM we detected in aggregates (Figure 1C-2E). Sugar-related proteins, whether in terms of metabolism (*e.g*. beta-galactosidase, glucanase), transfer (*e.g*. glycosyltransferase) or affinity (lectins), represent an important category of the candidates we identified in our multiomic analysis. Sugars could have a role in the ECM, or in modifying the properties of the cell wall which contains many glycoproteins ^43^. Glucan 1,3-Beta-Glucosidase of which 2 members are deregulated in *saz* mutants could be particularly interesting, since beta-glucans are glucose polymers that play a role in the response to stress in both plants ^44^ and animals ^45^. Finally, we analyzed a family of extracellular proteins, the VSPs (vegetative, SP rich) which are hydroxyproline and Serine-Proline (SP) rich proteins in Chlamydomonas enriched during the vegetative part of its cycle ^29^. Three VSP proteins (VSP4, VSP6 and VSP7) are downregulated in the culture medium of all *saz* mutants (Table S2). We further show here that an insertion in the VSP4 gene induces the formation of multicellular structures (Figure 6B) suggesting an anti-aggregative function for VSP4.

The protein content of Chlamydomonas medium has already been analyzed in the context of saline stress inducing the formation of palmelloids ^46^. Cross-analysis of these results with ours, shows that the formation of palmelloids and large aggregates involves specific sets of proteins. Indeed, even if MMPs or pherophorins are found in the secretome during palmelloid formation, their identities are distinct from those identified in the medium of aggregates. Moreover, among the 62 proteins identified during the formation or dislocation of palmelloids, only one (MMP3) is common to our list of 131 proteins of interest, showing that the formation of aggregates involves a protein network completely specific to this process.

Genes possibly involved in intracellular signaling during aggregation were suggested by our transcriptomic study. Several genes related to cGMP are inhibited in *saz* mutants, including five guanylate cyclase genes, which are responsible for its synthesis (Table S2). This suggests that this secondary messenger could participate in the signaling pathway leading to aggregation. Interestingly, in animals the destruction of ECM is mediated by MMPs that are activated by a signaling pathway involving cGMP ^47,48^. This activation occurs through the activation of kinases, which regulate gene expression, and may also involve the MAP kinase signaling pathway ^28^. Moreover, the two most represented families in our transcriptomic analysis are those of ECM-related proteins and protein kinases, among which we identified a MAP kinase kinase (MAP2K), whose gene is under-expressed in *saz* mutants. The fact that the signaling pathway known in animals to destroy the ECM through the activation of MMPs is inhibited in *saz* mutants is consistent with an important role of MMPs and ECM regulation for the establishment of multicellular structures in Chlamydomonas. We now plan to investigate the role of cGMP as a secondary messenger in the signaling pathway leading to aggregation.

In conclusion, we describe in this study the discovery of a new mode of adaptation of Chlamydomonas in response to stress, which involves the formation of multicellular structures that can comprise several thousand cells. We show that this process is fundamental, since within these aggregates the cells can survive environmental stresses. VLE seems to play a particularly important role in this process, since its absence leads to aggregation in *saz1* and it is downregulated in the extracellular compartment of four other *saz* mutants. We detected within the aggregates a sugar-rich ECM which, based on the results of our multiomics and reverse genetic analyses, appears to be central to the formation of the aggregates. We hypothesize that an intracellular signaling pathway partly involving the regulation of gene expression leads to an equilibrium in the extracellular compartment between pro- and anti-aggregative proteins such as pherophorins or MMPs, to control the establishment of protective multicellular structures. Comparison of the genomes of Chlamydomonas and *Volvox carteri* revealed a high degree of similarity between the gene families present, but among the protein families that diverged the most in Volvox appeared the MMPs and the pherophorins^10,49^. This led to the suggestion that the increase in the number and diversity of these proteins could be at the origin of the establishment of an ECM making the transition to multicellularity possible^9,50^. The results we present here could therefore constitute the missing link between stress response, ECM, aggregation and transition to multicellularity. Our discovery allows to explain how, during evolution, long periods of stress could have induced and stabilized the expression of MMPs and pherophorins, allowing the formation of aggregates containing an ECM, likely to have evolved, through the diversification of pherophorins, towards a stable structure such as the one known in *Volvox carteri*. These multicellular structures would have been even more advantageous during stress periods of evolution, since we have shown that they constitute a very efficient survival strategy. This scenario is all the more likely since the hypothesis classically formulated for the emergence of multicellularity, is the formation of an intermediate stage of aggregation^51^. The order of Volvocales is particularly admirably adapted to study this phenomenon, given that it extends over a relatively short evolutionary distance, from the unicellular Chlamydomonas to the multicellular *Volvox carteri*.

## Materials and Methods

### Strains, media, and growth condition

The D66 strain (CC-4425) ^12^, was used as a wild type for this study in addition to the wild-type strain of the CLiP library CMJ030 (CC-4533) ^22^, when necessary. The list of the CLiP mutants used in this study is shown in table S3. Cells were grown in liquid cultures mixotrophically in Tris acetate phosphate (TAP) medium ^52^ on a rotary shaker (120 rpm) under continuous light (40-60 μmol photons.m^−2^·s^−1^), at 25°C. Aggregation assay and stress treatments were done in 24-well plates in a 1 mL volume at a concentration of 4-8 × 10^6^ cells·mL^−1^. Rose bengal (330000) and paraquat (856177) were purchased from Sigma-Aldrich (Saint-Louis, USA). S-Nitrosoglutathione (GSNO) was synthesized as described in ^53^.

### Aggregate area and cell death image analysis

Multi-well plates were imaged with the Perfection V800 scanner (Epson, Suwa, Japan) and micrograph were taken using the Axio Observer (Zeiss, Oberkochen, Germany) microscope. The software used were Fiji / ImageJ 1.52a ^19^ and R Studio 1.2.5 (tidyverse, ggpubr and PlotsOfData) ^54,55^. The particle areas were determined from 24-well plate scans using a Fiji macro (github.com/fdecarpentier/ParticleWell). The quantification of cell death was done following the previously published method ^56^, using a final concentration of 0.2% w/v Evans Blue (E2129, Sigma-Aldrich). The percentage of dead cells was calculated on a minimum of 100 individuals. To quantify the area and viability of each particle, we created a Fiji macro (github.com/fdecarpentier/PlantDeath) based on the CIELAB color space.

### mVenus plasmid construction

Unless otherwise specified, the enzymes were bought from New England BioLabs (Ipswich, USA). The level one plasmid pCM1-034 was constructed by cloning the parts P_psad_ (pCM0-016), mVenus (pCM0-086) and T_psad_ (pCM0-114) ^57,58^ in the acceptor plasmid pICH47732 ^59^ by a Golden Gate reaction. pCM1-034 was then assembled with the blasticidin resistance transcription unit pCM1-029 ^56^ and the empty linker pICH50881 in the level M acceptor plasmid pAGM8031 ^59^ yielding pCMM-23.

### Chlamydomonas transformation and generation of *saz* mutants

i72 cassette conferring paromomycin resistance was excised from pSL72 plasmid ^60^ using *Xho*I and *Eco*RI, isolated on agarose gel and purified (Macherey-Nagel NucleoSpin Gel and PCR Clean-up Kit). Transformants were generated using electroporation as previously described ^61^, and selected on agar plates containing paromomycin at a concentration of 20 mg/L. The transformants were then grown in 96-well plates from which were selected the *saz* mutants that aggregated spontaneously.

For the generation of the mVenus strain, pCMM-23 was electroporated in D66 as previously described ^62^. Transformants were selected agar plates using with 50 mg/L blasticidin ^56^. After 6 days of growth, the plates were scanned with a Typhoon FLA 9500 laser scanner (GE, Healthcare) at 473 nm excitation with the filters BPG1 for YFP and LPG for chlorophyll, to select the mVenus fluorescent strain.

### Insert localization in *saz1* and CLiP mutants

Genomic DNA purification method was adapted from ^63^. The insertion locus of i72 cassette was determined by RESDA-PCR (Restriction Enzyme Site-Directed Amplification PCR) ^20^. PCR were performed with Quick-Load® *Taq* 2x Master Mix (New England BioLabs) according to the manufacturer recommendations with the primer displayed in Table S3. We used the NucleoSpin Gel and PCR Clean-up Kit (Macherey-Nagel) for band purification and sequencing was performed by Eurofins Genomics.

### Medium purification and protein quantitation by tandem mass spectrometry

Culture medium was harvested after 5 days of culture in TAP medium and cleared of cells by centrifugation (2300 × *g*, 5 min) and 0.45 μm filtration. For quantitative proteomics, 50 mL of the medium was concentrated 125 times using Amicon centrifugal filter unit (10 kDa MWCO, 4000 × *g*, 4°C). The concentrated medium was centrifugated (10 min, 17000 x *g*, 4°C) to pellet the insoluble debris. Twenty micrograms of proteins were denatured in the presence of Laemmli buffer and separated by SDS-PAGE. After a short migration (< 0.5 cm) and Coomassie blue staining, gel pieces containing extracellular proteins were excised and subjected to trypsin digestion as previously described (Marchand et al., 2010). Peptide mixtures were prepared in 60 μL of solvent A (0.1% (v/v) formic acid in 3% (v/v) acetonitrile). For each strain, five biological replicates were prepared which were hereafter analyzed as technical duplicates. Mass spectrometry analyses were performed on a Q-Exactive Plus hybrid quadripole-orbitrap mass spectrometer (Thermo Fisher, San José, CA, USA) coupled to an Easy 1000 reverse phase nano-flow LC system (Proxeon) using the Easy nano-electrospray ion source (Thermo Fisher). Five microliters of peptide mixtures were loaded onto an Acclaim PepMap™ precolumn (75 μm × 2 cm, 3 μm, 100 Å; Thermo Scientific) equilibrated in solvent A and separated at a constant flow rate of 250 nl/min on a PepMap™ RSLC C18 Easy-Spray column (75 μm × 50 cm, 2 μm, 100 Å; Thermo Scientific) with a 90 min gradient (0 to 20% B solvent (0.1% (v/v) formic acid in acetonitrile) in 70 min and 20 to 32% B solvent in 20 min). Data acquisition was performed as described in Pérez-Pérez et al., 2017. Raw Orbitrap data were processed with MaxQuant 1.5.6.5 using the Andromeda search engine ^66^ against the Chlamydomonas database (V.5.6) ^67,68^ and the MaxQuant contaminants database. Mass tolerance was set to 10 ppm for the parent ion mass and 20 mDa for fragments, and up to two missed cleavages per peptide were allowed. Peptides were identified and quantified using the “match between run” setting and an FDR (False Discovery Rate) below 0.01. The intensity of proteins with at least two unique peptides were quantified with the MaxLFQ method ^66^. Proteins detected in at least three biological replicates that were analyzed with a Wilcoxon-Mann-Whitney non parametrical test. Proteins were regarded as differentially expressed than the wild type for adjusted p-values < 0.05 and log_2_ (Fold Change) >1.

### RNA analysis

Total RNA of Chlamydomonas was extracted from 10 mL cultures at 5-6 × 10^6^ cells/mL according to the protocol described in Cavaiuolo et al., 2017. RNAs were treated with DNase I (New England BioLabs) according to the manufacturer recommendations. RNA quality was checked with the TapeStation System (Agilent, Santa Clara, USA), three biological replicates per strain with a RIN^e^ (RNA integrity number equivalent) above 5.5 were selected. Library preparation and sequencing were performed by the iGenSeq genotyping and sequencing core facility (ICM Institute - Hôpital Pitié-Salpêtrière AP-HP, Paris, France) using KAPA HyperPrep Kits (Roche, Basel, Switzerland) and the NovaSeq 6000 sequencer (Illumina, San Diego, USA). Paired-end reads were mapped against the Chlamydomonas genome v.5.6 using Bowtie2 v2.3.4.3 ^67,70^. with at least 30 base-mean mapped reads. Normalization and differential analysis were performed according to DESeq2 v1.30.0 ^71^. Genes were regarded as differentially expressed than the wild type for adjusted p-values < 0.05 and log_2_ (Fold Change) > 2 or < −2. For quantitative reverse transcription PCR analysis, RNA was treated with RNase-Free DNase (New England Biolabs) and reverse-transcribed following manufacturer’s recommendations (The ProtoScript Taq RT-PCR Kit, New England Biolabs). Relative mRNA, abundance was calculated using the comparative delta-Ct method, and normalized to the corresponding *RACK1* (Cre06.g278222) gene levels.

## Author contributions

A.D. and F.C. designed the study, A.D., F.C., and S.D.L. wrote the manuscript, A.D., F.C., A.M., C.M., C.C. and C.D. performed experiments and analyzed the data, P.C. assisted with modular cloning experiments.

## Acknowledgments

We acknowledge Marion Hamon of the mass spectrometry platform of the Institut de Biologie Physico-Chimique for the mass spectrometer running and data collection. We thank Oliver Caspari for his help in the fluorescent protein expression, and Lionel Bénard for his highly valuable advices about RNA extraction. Finally, we thank Nicolas D. Boisset, Théo Le Moigne, and Julien Henri for stimulating discussions and suggestions.

## References

1. Saur, I. M. L. Recognition and defence of plant-infecting fungal pathogens. Journal of Plant Physiology 14 (2021).

2. Flemming, H.-C. et al. Biofilms: an emergent form of bacterial life. Nat Rev Microbiol 14, 563–575 (2016).

3. Váchová, L. & Palková, Z. How structured yeast multicellular communities live, age and die? FEMS Yeast Research 18, (2018).

4. Vlamakis, H., Chai, Y., Beauregard, P., Losick, R. & Kolter, R. Sticking together: building a biofilm the Bacillus subtilis way. Nat Rev Microbiol 11, 157–168 (2013).

5. de Carpentier, F., Lemaire, S. D. & Danon, A. When Unity Is Strength: The Strategies Used by *Chlamydomonas* to Survive Environmental Stresses. Cells 8, 1307 (2019).

6. Sathe, S. & Durand, P. M. Cellular aggregation in *Chlamydomonas* (Chlorophyceae) is chimaeric and depends on traits like cell size and motility. European Journal of Phycology 51, 129–138 (2016).

7. Herron, M. D., Ratcliff, W. C., Boswell, J. & Rosenzweig, F. Genetics of a de novo origin of undifferentiated multicellularity. R Soc Open Sci 5, (2018).

8. Ratcliff, W. C. et al. Experimental evolution of an alternating uni- and multicellular life cycle in *Chlamydomonas reinhardtii*. Nature Communications 4, (2013).

9. Nishii, I. & Miller, S. M. Volvox: Simple steps to developmental complexity? Current Opinion in Plant Biology 13, 646–653 (2010).

10. Hanschen, E. R. et al. The Gonium pectorale genome demonstrates co-option of cell cycle regulation during the evolution of multicellularity. Nat Commun 7, 11370 (2016).

11. Kubo, T. et al. The Chlamydomonas Hatching Enzyme, Sporangin, is Expressed in Specific Phases of the Cell Cycle and is Localized to the Flagella of Daughter Cells Within the Sporangial Cell Wall. Plant and Cell Physiology 50, 572–583 (2009).

12. Schnell, R. A. & Lefebvre, P. A. Isolation of the Chlamydomonas Regulatory Gene *NIT2* by Transposon Tagging. Genetics 134, 737–747 (1993).

13. Morisse, S., Zaffagnini, M., Gao, X.-H., Lemaire, S. D. & Marchand, C. H. Insight into Protein S-nitrosylation in *Chlamydomonas reinhardtii*. Antioxidants & Redox Signaling 21, 1271–1284 (2014).

14. Fischer, B. B., Krieger-Liszkay, A. & Eggen, R. I. L. Oxidative stress induced by the photosensitizers neutral red (type I) or rose bengal (type II) in the light causes different molecular responses in *Chlamydomonas reinhardtii*. Plant Science 168, 747–759 (2005).

15. Laloi, C. et al. Cross-talk between singlet oxygen- and hydrogen peroxide-dependent signaling of stress responses in Arabidopsis thaliana. Proc. Natl. Acad. Sci. U.S.A. 104, 672–677 (2007).

16. Schroda, M., Hemme, D. & Mühlhaus, T. The Chlamydomonas heat stress response. Plant J 82, 466–480 (2015).

17. Cavada, B., Pinto-Junior, V., Osterne, V. & Nascimento, K. ConA-Like Lectins: High Similarity Proteins as Models to Study Structure/Biological Activities Relationships. IJMS 20, 30 (2018).

18. Preethi, N., Vanitha, P., Ramu, V., Sheshshayee, M. & Udayakumar, M. Quantification of Membrane Damage/Cell Death Using Evan’s Blue Staining Technique. BIO-PROTOCOL 7, (2017).

19. Schindelin, J. et al. Fiji - an Open Source platform for biological image analysis. Nat Methods 9, (2012).

20. González-Ballester, D., de Montaigu, A., Galván, A. & Fernández, E. Restriction enzyme site-directed amplification PCR: A tool to identify regions flanking a marker DNA. Analytical Biochemistry 340, 330–335 (2005).

21. Matsuda, Y., Koseki, M., Shimada, T. & Saito, T. Purification and characterization of a vegetative lytic enzyme responsible for liberation of daughter cells during the proliferation of Chlamydomonas reinhardtii. Plant Cell Physiol. 36, 681–689 (1995).

22. Li, X. et al. A genome-wide algal mutant library and functional screen identifies genes required for eukaryotic photosynthesis. Nat Genet 51, 627–635 (2019).

23. Tardif, M. et al. PredAlgo: A New Subcellular Localization Prediction Tool Dedicated to Green Algae. Molecular Biology and Evolution 29, 3625–3639 (2012).

24. Hallmann, A. The pherophorins: common, versatile building blocks in the evolution of extracellular matrix architecture in Volvocales. The Plant Journal 45, 292–307 (2006).

25. Abe, J. et al. The transcriptional program of synchronous gametogenesis in Chlamydomonas reinhardtii. Curr Genet 46, 304–315 (2004).

26. Kinoshita, T., Fukuzawa, H., Shimada, T., Saito, T. & Matsuda, Y. Primary structure and expression of a gamete lytic enzyme in Chlamydomonas reinhardtii: similarity of functional domains to matrix metalloproteases. Proceedings of the National Academy of Sciences 89, 4693–4697 (1992).

27. Nishimura, M., Nagashio, R., Sato, Y. & Hasegawa, T. Late Somatic Gene 2 disrupts parental spheroids cooperatively with Volvox hatching enzyme A in Volvox. Planta 245, 183–192 (2017).

28. Zaragoza, C. et al. Activation of the Mitogen Activated Protein Kinase Extracellular Signal-Regulated Kinase 1 and 2 by the Nitric Oxide–cGMP–cGMP–Dependent Protein Kinase Axis Regulates the Expression of Matrix Metalloproteinase 13 in Vascular Endothelial Cells. Mol Pharmacol 62, 927–935 (2002).

29. Waffenschmidt, S., Woessner, J. P., Beer, K. & Goodenough, U. W. Isodityrosine cross-linking mediates insolubilization of cell walls in Chlamydomonas. Plant Cell 5, 809 (1993).

30. Laczko, R. & Csiszar, K. Lysyl Oxidase (LOX): Functional Contributions to Signaling Pathways. Biomolecules 10, 1093 (2020).

31. Alieva, N. O. et al. Myosin IIA and formin dependent mechanosensitivity of filopodia adhesion. Nat Commun 10, 3593 (2019).

32. Eisenmann, K. M. et al. T Cell Responses in Mammalian Diaphanous-related Formin mDia1 Knock-out Mice. J. Biol. Chem. 282, 25152–25158 (2007).

33. Niethamer, T. K. & Bush, J. O. Getting direction(s): The Eph/ephrin signaling system in cell positioning. Dev. Biol. 447, 42–57 (2019).

34. Liu, P. et al. Chlamydomonas PKD2 organizes mastigonemes, hair-like glycoprotein polymers on cilia. Journal of Cell Biology 219, e202001122 (2020).

35. Huang, K. et al. Function and dynamics of PKD2 in Chlamydomonas reinhardtii flagella. Journal of Cell Biology 179, 501–514 (2007).

36. Bernardes, J. P. et al. The evolution of convex trade-offs enables the transition towards multicellularity. Nat Commun 12, 4222 (2021).

37. Kim, C. & Sharma, R. Cyclic nucleotide selectivity of protein kinase G isozymes. Protein Science 30, 316–327 (2021).

38. Luxmi, R. et al. Proteases Shape the Chlamydomonas Secretome: Comparison to Classical Neuropeptide Processing Machinery. Proteomes 6, 36 (2018).

39. Snell, W. J., Eskue, W. A. & Buchanan, M. J. Regulated secretion of a serine protease that activates an extracellular matrix-degrading metalloprotease during fertilization in Chlamydomonas. Journal of Cell Biology 109, 1689–1694 (1989).

40. Marino, G. & Funk, C. Matrix metalloproteinases in plants: a brief overview. Physiologia Plantarum 145, 196–202 (2012).

41. Fanjul-Fernández, M., Folgueras, A. R., Cabrera, S. & López-Otín, C. Matrix metalloproteinases: Evolution, gene regulation and functional analysis in mouse models. Biochimica et Biophysica Acta (BBA) - Molecular Cell Research 1803, 3–19 (2010).

42. Sumper, M., Berg, E., Wenzl, S. & Godl, K. How a sex pheromone might act at a concentration below 10(−16) M. The EMBO Journal 12, 831–836 (1993).

43. Voigt, J. & Frank, R. 14-3-3 Proteins Are Constituents of the Insoluble Glycoprotein Framework of the Chlamydomonas Cell Wall. Plant Cell 15, 1399–1413 (2003).

44. Fesel, P. H. & Zuccaro, A. β-glucan: Crucial component of the fungal cell wall and elusive MAMP in plants. Fungal Genetics and Biology 90, 53–60 (2016).

45. Hopke, A., Brown, A. J. P., Hall, R. A. & Wheeler, R. T. Dynamic fungal cell wall architecture in stress adaptation and immune evasion. 19 (2019).

46. Khona, D. K. et al. Characterization of salt stress-induced palmelloids in the green alga, Chlamydomonas reinhardtii. Algal Research 16, 434–448 (2016).

47. Denninger, J. W. & Marletta, M. A. Guanylate cyclase and the cNO/cGMP signaling pathway. Biochimica et Biophysica Acta 334–350 (1999).

48. Dey, N. B. & Lincoln, T. M. Possible involvement of Cyclic-GMP-dependent protein kinase on matrix metalloproteinase-2 expression in rat aortic smooth muscle cells. Mol Cell Biochem 368, 27–35 (2012).

49. Prochnik, S. E. et al. Genomic Analysis of Organismal Complexity in the Multicellular Green Alga Volvox carteri. Science 329, 223–226 (2010).

50. Olson, B. J. & Nedelcu, A. M. Co-option during the evolution of multicellular and developmental complexity in the volvocine green algae. Current Opinion in Genetics & Development 39, 107–115 (2016).

51. Olson, B. J. From brief encounters to lifelong unions. eLife 2, e01893 (2013).

52. Gorman, D. S. & Levine, R. P. Cytochrome f and plastocyanin: their sequence in the photosynthetic electron transport chain of Chlamydomonas reinhardi. Proc Natl Acad Sci U S A 54, 1665–1669 (1965).

53. Tagliani, A. et al. Structural and functional insights into nitrosoglutathione reductase from Chlamydomonas reinhardtii. Redox Biology 38, 101806 (2021).

54. Postma, M. & Goedhart, J. PlotsOfData—A web app for visualizing data together with their summaries. PLOS Biology 17, e3000202 (2019).

55. Wickham, H. ggplot2: Elegant Graphics for Data Analysis. (Springer, 2016).

56. de Carpentier, F. et al. Blasticidin S Deaminase: A New Efficient Selectable Marker for *Chlamydomonas reinhardtii*. Front. Plant Sci. 11, (2020).

57. Crozet, P. et al. Birth of a Photosynthetic Chassis: A MoClo Toolkit Enabling Synthetic Biology in the Microalga *Chlamydomonas reinhardtii*. ACS Synth. Biol. 7, 2074–2086 (2018).

58. Lauersen, K. J., Kruse, O. & Mussgnug, J. H. Targeted expression of nuclear transgenes in *Chlamydomonas reinhardtii* with a versatile, modular vector toolkit. Appl Microbiol Biotechnol 99, 3491–3503 (2015).

59. Weber, E., Engler, C., Gruetzner, R., Werner, S. & Marillonnet, S. A Modular Cloning System for Standardized Assembly of Multigene Constructs. PLoS ONE 6, e16765 (2011).

60. Pollock, S. V., Prout, D. L., Godfrey, A. C., Lemaire, S. D. & Moroney, J. V. The *Chlamydomonas reinhardtii* proteins Ccp1 and Ccp2 are required for long-term growth, but are not necessary for efficient photosynthesis, in a low-CO2 environment. Plant Mol Biol 56, 125–132 (2004).

61. Pollock, S. V. et al. A robust protocol for efficient generation, and genomic characterization of insertional mutants of *Chlamydomonas reinhardtii*. Plant Methods 13, 22 (2017).

62. Onishi, M. & Pringle, J. R. Robust Transgene Expression from Bicistronic mRNA in the Green Alga *Chlamydomonas reinhardtii*. G3: Genes, Genomes, Genetics 6, 4115–4125 (2016).

63. Danon, A. & Gallois, P. UV-C radiation induces apoptotic-like changes in *Arabidopsis thaliana*. FEBS Letters 437, 131–136 (1998).

64. Marchand, C. H. et al. Thioredoxin targets in *Arabidopsis* roots. PROTEOMICS 10, 2418–2428 (2010).

65. Pérez-Pérez, M. E. et al. The Deep Thioredoxome in *Chlamydomonas reinhardtii:* New Insights into Redox Regulation. Molecular Plant 10, 1107–1125 (2017).

66. Tyanova, S., Temu, T. & Cox, J. The MaxQuant computational platform for mass spectrometry-based shotgun proteomics. Nature Protocols 11, 2301–2319 (2016).

67. Blaby, I. K. et al. The Chlamydomonas genome project: a decade on. Trends in Plant Science 19, 672–680 (2014).

68. Gallaher, S. D. et al. High-throughput sequencing of the chloroplast and mitochondrion of *Chlamydomonas reinhardtii* to generate improved *de novo* assemblies, analyze expression patterns and transcript speciation, and evaluate diversity among laboratory strains and wild isolates. The Plant Journal 93, 545–565 (2018).

69. Cavaiuolo, M., Kuras, R., Wollman, F.-A., Choquet, Y. & Vallon, O. Small RNA profiling in *Chlamydomonas:* insights into chloroplast RNA metabolism. Nucleic Acids Res 45, 10783–10799 (2017).

70. Langmead, B. & Salzberg, S. L. Fast gapped-read alignment with Bowtie 2. Nature Methods 9, 357–359 (2012).

71. Love, M. I., Huber, W. & Anders, S. Moderated estimation of fold change and dispersion for RNA-seq data with DESeq2. Genome Biology 15, 550 (2014).

