## Supplementary material for "Stress-induced collective behavior leads to the formation of multicellular structures and the survival of the unicellular alga Chlamydomonas": Figure S1

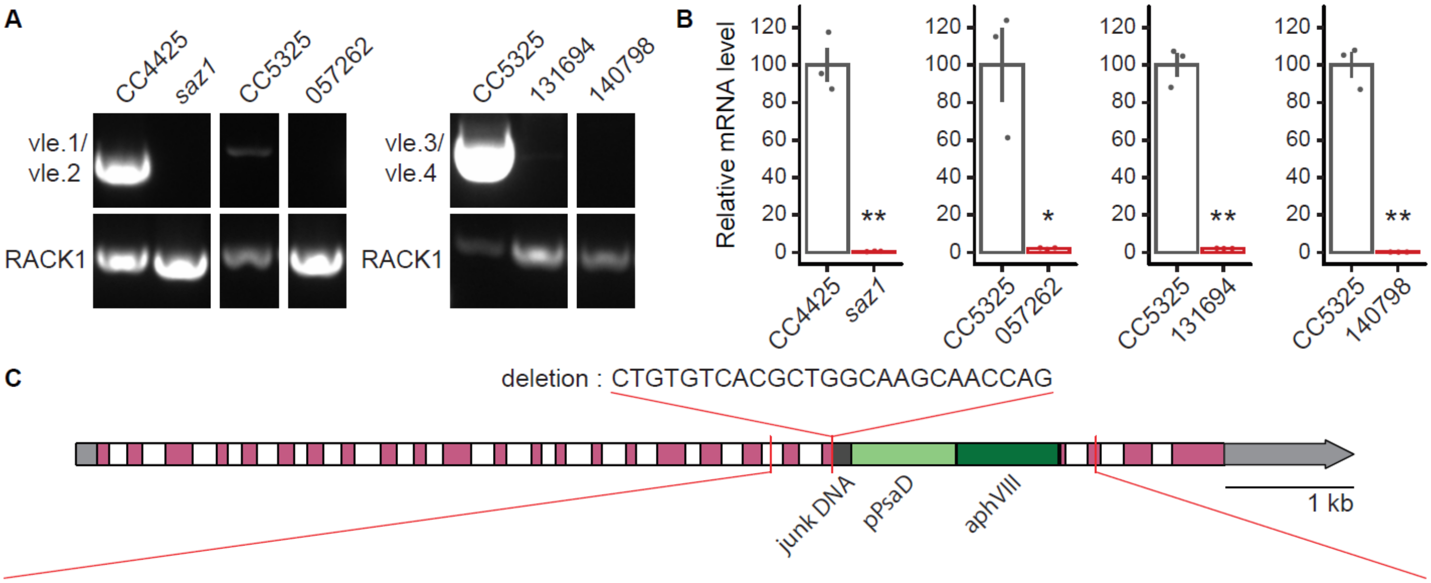


5'-acgcggccacaagctgcagctcaagggtgtgttaccgtagtcccgtgcattaaggccatgctccattcgctttttgcttttcccaccttgcagCTTCGAAGCGGTCTCTAT
GGCACCGGCACCGCCATGCTCAACCCCGCCAACTATGTGGTGTATGGCTATGCCCGCACCAACCTTGCGCCGAGGGCCTTC
AGCCAGGGCGTGGCCTTCCGCCTGCGCgtgagtgcaaaatgtccgcgccgcacacatcttggcctttttaaaagagaaaccgatgtaaaggcgcttgcttgtggc
cactgttatgcaaagcctctttctgaatacctgtcaatatgtcccacttgtagctgcgcttagcgttcctcgtgcgcgtgtgctgcttctcaaccgaccacagGGCCCGCACACGC
GCCTGTTCATCAACGGGCAGTTGGTCTATGACTTCCAGTCCACGTCCGTGTTCCAGACCGTCACGCCTTGCCGCGTGTGGC
CCAGCTGGTCCTGGCCGCCAGCAATCAGTTCTCTGAGCAGATGTCGGCCACCAACGGCAACCGGTTCGCGTGGCTGCAGG
ATAGCCAGGCCGGCGACAGCAAGGCTGGCTGTGTTCCAGCGGTTCCACGGTTCAGCACACACCTGCCCGTCTGCCTGACA
GGAAGTGAACGCATGTCGAGGGAGGCCTCACCAATCGTCACACGAGCCCTCGTCAGAAACACGTCTCCGCCACGCTCTCC
CTCTCACGGCCGACCCCGCAGCCCTTTTGCCCTTTCCTAGGCCACCGACAGGACCCAGGCGCTCTCAGCATGCCTCAACAA
CCCGTACTCGTGCCAGCGGTGCCCTTGTGCTGGTGATCGCTTGGAAGCGCATGCGAAGACGAAGGGGCGGAGCAGGCGG
CCTGGCTGTTCGAAGGGCTCGCCGCCAGTTCGGGTGCCTTTCTCCACGCGCGCCTCCACACCTACCGATGCGTGAAGGCA
GGCAAATGCTCATGTTTGCCCGAACTCGGAGTCCTTAAAAAGCCGCTTCTTGTCGTCGTTCCGAGACATGTTAGCAGATCGC
AGTGCCACCTTTCCTGACGCGCTCGGCCCCATATTCGGACGCAATTGTCATTTGTAGCACAATTGGAGCAAATCTGGCGAGG
CAGTAGGCTTTTAAGTTGCAAGGCGAGAGAGCAAAGTGGGACGCGGCGTGATTATTGGTATTTACGCGACGGCCCGGCGCG
TTAGCGGCCCTTCCCCCAGGCCAGGGACGATTATGTATCAATATTGTTGCGTTCGGGCACTCGTGCGAGGGCTCCTGCGGG
CTGGGGAGGGGGATCTGGGAATTGGAGGTACGACCGAGATGGCTTGCTCGGGGGGAGGTTTCCTCGCCGAGCAAGCCAG
GGTTAGGTGTTGCGCTCTTGACTCGTTGTGCATTCTAGGACCCCACTGCTACTCACAACAAGCCCATATGGACGATGCGTTG
CGTGCACTGCGGGGTCGGTATCCCGGTTGTGAGTGGGTTGTTGTGGAGGATGGGGCCTCGGGGGCTGGTGTTTATCGGCT
TCGGGGTGGTGGGCGGGAGTTGTTTGTCAAGGTGGCAGCTCTGGGGGCCGGGGTGGGCTTGTTGGGTGAGGCTGAGCG
GCTGGTGTGGTTGGCGGAGGTGGGGATTCCCGTACCTCGTGTTGTGGAGGGTGGTGGGGACGAGAGGGTCGCCTGGTTG
GTCACCGAAGCGGTTCCGGGGCGTCCGGCCAGTGCGCGGTGGCCGCGGGAGCAGCGGCTGGACGTGGCGGTGGCGCT
CGCGGGGCTCGCTCGTTCGCTGCACGCGCTGGACTGGGAGCGGTGTCCGTTCGATCGCAGTCTCGCGGTGACGGTGCCG
CAGGCGGCCCGTGCTGTCGCTGAAGGGAGCGTCGACTTGGAGGATCTGGACGAGGAGCGGAAGGGGTGGTCGGGGGAG
CGGCTTCTCGCCGAGCTGGAGCGGACTCGGCCTGCGGACGAGGATCTGGCGGTTTGCCACGGTCACCTGTGCCCGGACA
ACGTGCTGCTCGACCCTCGTACCTGCGAGGTGACCGGGCTGATCGACGTGGGGCGGGTCGGCCGTGCGGACCGGCACTC
CGATCTCGCGCTGGTGCTGCGCGAGCTGGCCCACGAGGAGGACCCGTGGTTCGGGCCGGAGTGTTCCGCGGCGTTCCTG
CGGGAGTACGGGCGCGGGTGGGATGGGGCGGTATCGGAGGAAAAGCTGGCGTTTTACCGGCTGTTGGACGAGTTCTTCTG
AGATATCGAATTCTCGCACGAGATCTACCTCTACTTCGCTGCAATGgtgcgtgatgcgacgcaacgcagcaaggcagactgccacggccgtgggctt
cttggggtcgtgcagggcattgttgctacgtgctcagtgggcctcatcatgctgccaggtcgtagcccatgcgcatgtaaccctgcgtttgagcccggttgccctgtccccgcgcccttcct
gcagCTCAACAACACCGCGCCTGTGGGCATTCAGTGGGCCAACTGCAGCTCGCCCAACCAGCCCGCG-3'

Features :
5' flanking sequence : [5410 : 5889]
junk DNA : [5890 : 6037]
Cre07.g350600 : [5916 : 6006]
Cre16.g671250 : [6007 : 6025]
i72 : [6038 : 7671]
pPSAD : [6039 : 6852]
AphVIII : [6859 : 7656]
3' flanking sequence : [7672 : 7944]

Figure S1. Characterization of the different insertional mutants for the VLE gene. (A) For each of the mutants we verified by PCR that the insertion was in the predicted location in the VLE gene, using primers located on both sides of the insertion. Amplifications were detected only in the wild-type strains (CC4425 and CC5325) at the expected sizes, while in the mutants, amplification was made impossible due to the presence of the insertion. RACK1 (Cre06.g278222) was used as a positive control for the PCR. (B) Quantification of VLE gene expression by quantitative PCR in the saz1 mutant and the 3 other vle CLiP mutants20(Li et al., 2019)(Li et al., 2019). We used the primers vle2/vle5 to detect the insertion in saz1, vle1/vle6 for LMJ.RY0402.057262, vle7/vle8 for LMJ.RY0402.131694 and vle9/vle10 for LMJ.RY0402.140798, RACK1 was used as a positive control for the PCR. Values are mean of 3 biological replicates with 2 technical replicates. Error bars indicates ± SEM and symbols correspond to t-test p-values of * ≤ 0.05 or ** ≤ 10-2. (C) Sequencing of the cassette insertion zone in the saz1 mutant with primers from the gene and the cassette. Primer sequences can be found in Table S3.
