## Supplementary material for "Stress-induced collective behavior leads to the formation of multicellular structures and the survival of the unicellular alga Chlamydomonas": Figure S2

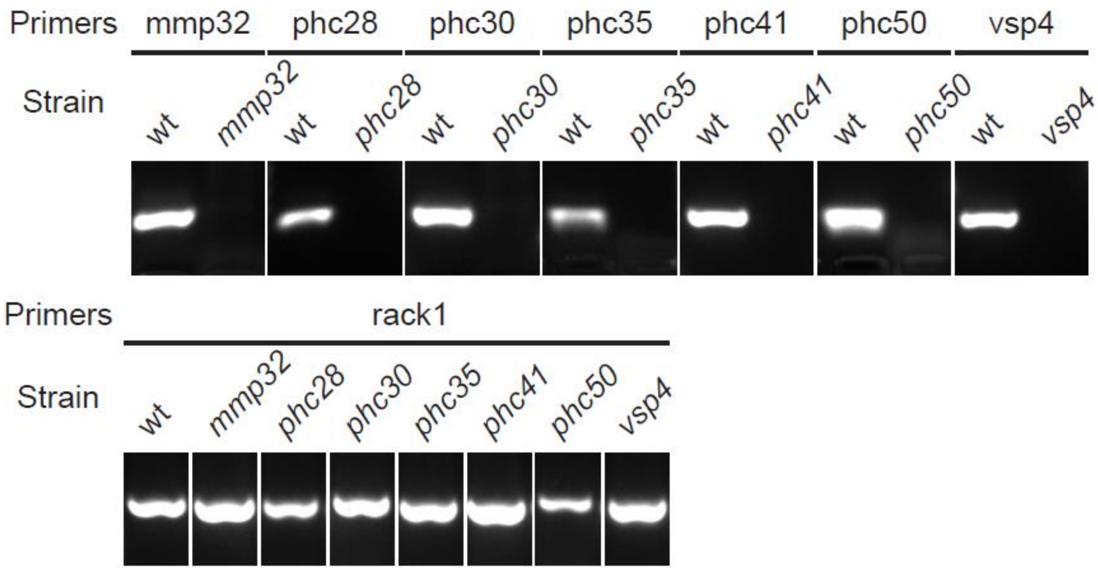


Figure S2. Genotyping of the CLiP mutants used in this study. PCR products obtained in the wild-type and the different mutants strains using control primers (RACK1), or primers located on both side of the insertion of the different mutants. The absence of PCR amplification in the mutants is due to the presence of the insert. Primers sequences and CLiP mutant names can be found in Table S3.
