## Supplementary material for "Stress-induced collective behavior leads to the formation of multicellular structures and the survival of the unicellular alga Chlamydomonas": Table S3

| target gene | Gene name | mutant | Primer name | Primer sequence |
|---|---|---|---|---|
| Cre01.g049950 | VLE | *saz1* | vle.1 | CCTCCAACATGCAAATCGG |
| | | LMJ.RY0402.057262 | vle.2 | CGGAAGAGAAGGACACGTAG |
| | | LMJ.RY0402.131694 | vle.3 | TGTCAGTGAATCTTGCTGGC |
| | | LMJ.RY0402.140798 | vle.4 | CCTGTTCTGTTTTGGCCAGT |
| | | *saz1* | vle.2 | CGGAAGAGAAGGACACGTAG |
| | | | vle.5 | CGTGGATATACCAGTACGGC |
| | | LMJ.RY0402.057262 | vle.1 | CCTCCAACATGCAAATCGG |
| | | | vle.6 | CGAAGTTGAAGTCAGGCACA |
| | | LMJ.RY0402.131694 | vle.7 | CCTGGAAAGGAGCGACGCAG |
| | | | vle.8 | TCTGCACCCGTGATGTGTAT |
| | | LMJ.RY0402.140798 | vle.9 | TGAATGACCCGCAACTGAGC |
| | | | vle.10 | AGCGAATGTGTTTTTGCCGATG |
| Cre14.g625850 | MMP32 | LMJ.RY0402.170721 | mmp32.1 | GCAGGTTCAGATTGTTTGTTGG |
| | | | mmp32.2 | AGCATGACGATGGATGTGATG |
| Cre17.g717950 | PHC28 | LMJ.RY0402.116054 | phc28.1 | AGTCAACTTCAATAGCTTCCG |
| | | | phc28.2 | TGTTCATCCATTATCTATGGCG |
| Cre09.g404201 | PHC30 | LMJ.RY0402.246644 | phc30.1 | ACAGTTGGTGGGGTTTGAG |
| | | | phc30.2 | TGCGGTTGTGATAGGTAGAAG |
| Cre05.g238687 | PHC35 | LMJ.RY0402.201505 | phc35.1 | CGTGAAGTTCTCGATCATTCC |
| | | | phc35.2 | TCGCCACAAATTGTCATGTAG |
| Cre12.g506750 | PHC41 | LMJ.RY0402.152127 | phc41.1 | ACCTCCACTTCTCGGTATG |
| | | | phc41.2 | TTCAGAACGTGTGATGACG |
| Cre06.g292249 | PHC50 | LMJ.RY0402.121429 | phc50.1 | TCTTCACAGCACCATCGCTC |
| | | | phc50.2 | ACAACCCACGTAGAAGTCGC |
| Cre09.g391801 | VSP4 | LMJ.RY0402.176992 | vsp4.1 | GCACTCGTCGTTACGCAATG |
| | | | vsp4.2 | CAAGTGTGGAAACGATTCGCC |
| i72 | i72 | NA | ppsad.1 | GCTGGCACGAGTACGGGTTG |
| | | | RB4 | TGGTTCGGGCCGGAGTGTTC |
| Cre06.g278222 | RACK1 | NA | rack1.5 | GACGTCATCCACTGCCTGTG |
| | | | rack1.3 | CGACGCATCCTCAACACACC |

Table S3. List of primers used in our study.

The genes we studied are referenced in this table as well as the CliP mutants and the corresponding primer pairs used.
